# Proteomic profiling of isolated immune synapses from primary mouse B cells

**DOI:** 10.1101/2023.02.23.529674

**Authors:** Diogo M. Cunha, Sara Hernández-Pérez, Luqman O. Awoniyi, Alexandre F. Carisey, Guillaume Jacquemet, Pieta K. Mattila

## Abstract

The immune synapse (IS) is a cell-cell interaction platform critical in lymphocyte activation by specific antigens. Despite of B cells being able to also respond to soluble antigens, in particular the *in vivo* importance of the IS and surface-tethered antigen recognition has strongly emerged in the recent years. The IS serves as a dynamic hub for multiple cellular actions but the molecular details of these functions, especially beyond the B cell antigen receptor (BCR) signalling, remain poorly understood. Here, to address the lack in the systems level understanding of the IS, we setup methodology for comprehensive investigation of the composition of the primary mouse B cells’ IS at proteome level. Utilizing functionalized magnetic beads to mimic antigen presenting cells and trigger IS formation on them, we developed a method to specifically and robustly extract the cell adhesions on the beads, namely the IS or transferrin receptor mediated adhesion as a control. Our data revealed 661 proteins exclusively present in the IS at 15 minutes after BCR engagement, 13 exclusively in the control adhesions and 365 proteins shared between the samples. We got strong coverage of the known components of the IS as well as identified a plethora of unknown proteins and functional pathways with hitherto unknown roles in B cell IS. Thus, in this work, we validated the IS isolation method as a valuable tool to study early B cell activation by surface-bound antigens as well as unveil several novel proteins and pathways suggestive of new functional aspects in the IS.

## Introduction

B lymphocytes are a vital part of the adaptive immunity as they produce specific, high-affinity antibodies against pathogens and orchestrate other branches of the immune response. To do so, B cells rely on specific antigen recognition by the B cell receptor (BCR). Although B cells strongly respond to soluble antigens, *in vivo* antibody responses are suggested to be more commonly triggered by the binding of foreign, unprocessed antigens on the surface of antigen-presenting cells (APCs) (Batista & Harwood, 2009; Kuokkanen et al., 2015; Shaheen et al., 2019). Different cells can act as APCs for B cells, including subcapsular sinus macrophages (Carrasco & Batista, 2007; Iliopoulou et al., 2022), dendritic cells, and follicular dendritic cells (Heath et al., 2019; Martinez-Riano et al., 2022; Qi et al., 2006; Suzuki et al., 2009). Upon antigen recognition, the interaction between the APC and the B lymphocyte triggers the formation of an activation platform termed immune synapse (IS) (Batista & Harwood, 2009), that serves as critical hub for cellular events linked to BCR signalling and lowers the threshold for receptor activation (Carrasco & Batista, 2007, Natkanski et al., 2013).

Antigen recognition by the BCR prompts a robust cascade of events, such as BCR signalling, BCR microcluster (or BCR signalosome) assembly (Geahlen, 2009; Marshall et al., 2018; Nicholson, 2016; Tedford et al., 2001) and contraction of the microclusters into the centre of the synapse for internalisation and further processing (Avalos & Ploegh, 2014; Hou et al., 2006). Thus, the IS plays an essential role in B cell activation, as it is not only a platform for signalling but also for physical sensing of the antigen properties and microenvironment, antigen acquisition and degradation (del Valle Batalla et al., 2018, Spillane and Tolar, 2018). To fulfil its functions, the IS encompasses various cell biological machineries that coordinate the IS formation and the tightly linked events, like vesicle traffic. One of the major players at the IS is the highly plastic actin cytoskeleton, that provides the force and support to the IS structure driving the cell spreading and contraction, as well as controls vesicle trafficking together with microtubule cytoskeleton. The actin cytoskeleton even controls the diffusion of the BCR and its co-receptors, CD19 and CD81, in the membrane influencing the signalling activity (Mattila et al., 2013, 2016; Treanor et al., 2010). Consequently, the formation of the IS requires such an extent of cellular processes that only a handful of techniques provide a solid approach to fully capture this complexity.

In the last decade, we have witnessed a vertiginous development of techniques, including advanced microscopy, genome-wide screens, next-generation sequencing (NGS) and mass spectrometry (MS)-based proteomics. These techniques have allowed for the identification of several proteins and mechanisms regulating BCR activation (Awoniyi et al.; Li et al., 2014; Malinova et al., 2020; Matsumoto et al., 2009; Satpathy et al., 2015). However, largely due to the easier methodology, the studies have mainly focused on the changes provoked after activation with soluble antigen, instead of the more *in vivo* relevant surface-bound antigen.

In this study, we developed a pull-down approach to extract the proteins forming the IS or dynamically recruited to the IS during mouse primary B cell activation. We used antigen-coated paramagnetic beads to mimic activation by surface-tethered antigen in APCs and to trigger IS formation. We utilized chemical crosslinking to stabilize the IS proteome, prior to sonication to remove cell bodies, to recover the bead-bound protein networks for MS analysis. As a control, we triggered adhesions by engagement of Transferrin receptor (TfR). With this approach, we identified 1026 proteins in the primary B cell IS, out of which 723 either exclusively present or significantly enriched in the IS compared to the control adhesions. The resulting proteome largely covered the well-known components of BCR signalosome but also identified a multitude of proteins not previously linked to IS function.

## Results

### The use of antigen-coated beads, but not anti-TfR-coated beads, efficiently triggers B cell activation

In order to isolate B cell IS for proteomic analysis, inspired by the protocols developed for the study of integrin adhesions (Byron et al., 2015; Jones et al., 2015), we set-up a method utilizing functionalized paramagnetic beads in conjunction with chemical cross-linking and removal of cell bodies by sonication to enable specific and comprehensive pull-down of cellular adhesions (Fig.1A, schematics).

**Figure 1.**
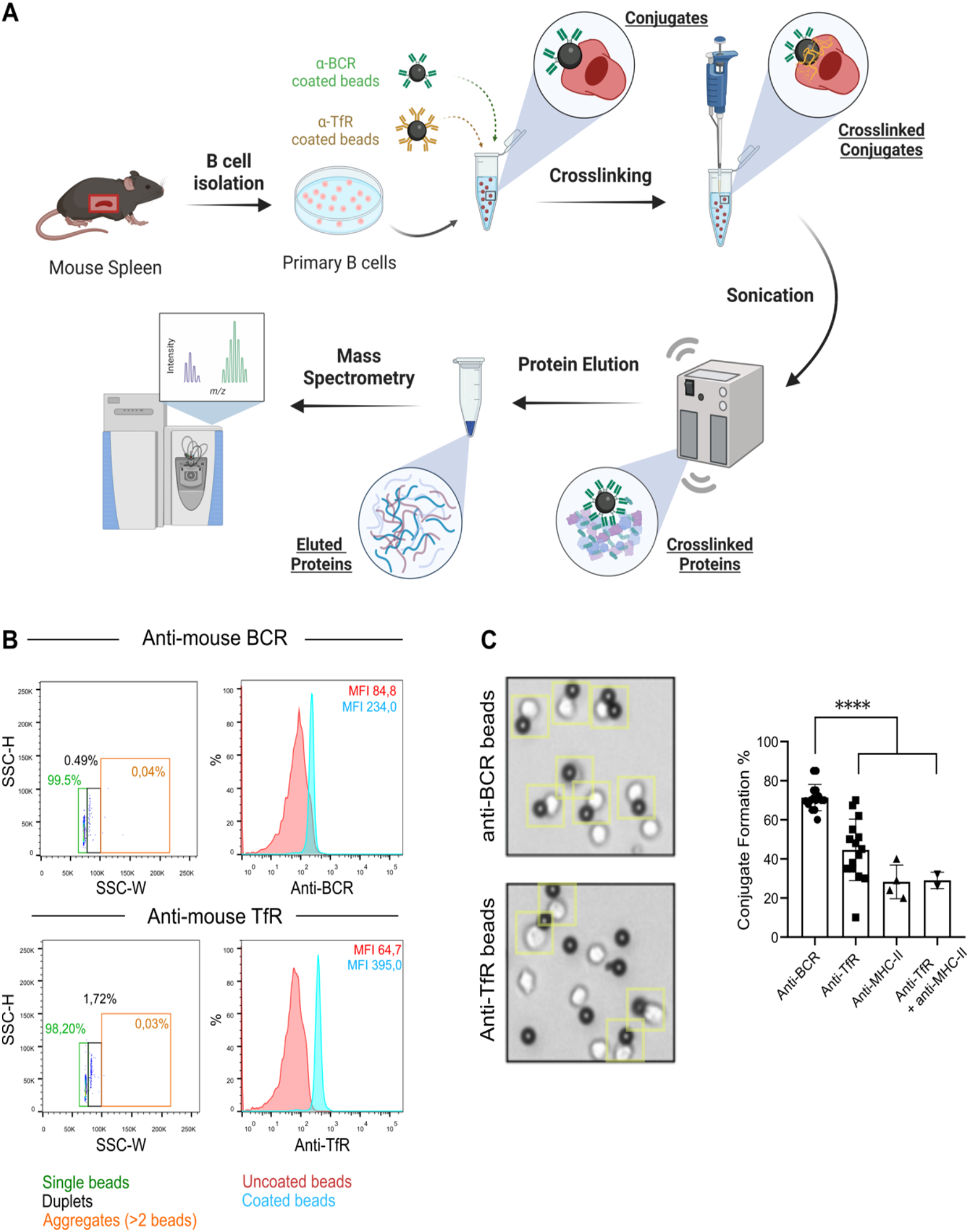
The isolation of the primary B cell IS. (A) Schematic representation of the IS isolation method. Primary mouse B cells are let to form the IS on magnetic beads and the IS structures are extracted for proteomic analysis. (B) Coating efficiency of the beads with selected antibodies was determined using flow cytometry. Single beads (green), duplets (black) and aggregates (orange) were gated separately (on the left) and the fluorescence intensity of the single beads was analysed (on the right). Red histogram: uncoated beads. Blue histogram: coated beads. (C) Cell-bead conjugate formation using beads coated with anti-BCR, anti-TfR, anti-MHC-II, or anti-TfR + anti-MHC-II antibodies. Primary cells were incubated with coated beads for 15 min and analysed using an inverted brightfield microscopy (on the left). Yellow squares highlight single conjugates. After conjugation, unbound cells were counted to get the percentage of conjugate formation (on the right). In the graph, the average percentage of conjugates [100 – (unbound cells/total cells)*100] per experiment are shown, from 2-15 experiments. Anti-BCR: 71,4 ± 1.7 % (*n* = 15). Anti-TfR 44.6 ± 4.0 % (*n* = 15). Anti-MHC-II: 28.3 ± 4.4 % (*n* = 4). Anti-CD71 + anti-MHC-II: 29.0 ± 3.0 % (*n* = 2). Each symbol represents and individual experiment. **** P < 0.0001 (unpaired Student’s t-test).

In order to mimic the antigens presented on the surface of APCs and to trigger IS formation, we employed F(ab’)2 anti-IgM-coated beads (anti-BCR beads from now on) as surrogate antigen, compatible with the use of mouse primary B cells. The use of such surface-bound surrogate antigen to activate B cells and study IS formation has been widely employed over the years, and it is thus well validated and characterised in a number of systems (e.g., beads, plastic plates, glass coverslips, supported-lipid bilayers, and acrylamide gels). However, there is lack of consensus regarding the use of a control ligand that triggers cell adhesion without activating BCR signalling. Several ligands have been used in the literature for this purpose, including, but not limited to, fibronectin, transferrin (Tf), anti-TfR, and anti-MHC-II antibodies (Ketchum et al., 2018; Roche & Furuta, 2015; Stupack et al., 1991). Thus, in addition to the anti-BCR, we tested these four ligands, to determine the most suitable control for our system. We examined 1) the efficiency of these ligands to coat the beads and 2) the ability of the primary B cells to form conjugates with the ligand-coated beads.

As shown by flow cytometry analysis, anti-BCR beads were effectively coated and essentially all of the beads were detected as single beads (Fig. 1B). Also anti-TfR beads (Fig. 1B), anti-MHCII beads (Supp. Fig. S1A), and a combination of anti-TfR and anti-MHC-II (Supp. Fig. 1B) showed successful coating without aggregation. Despite being directly fluorescently labelled, transferrin was not detected on the beads, suggesting that the coating was inefficient (Supp. Fig. S1C). Fibrinogen, on the other hand, exhibited very efficient coating, but also a high tendency to generate aggregates, as only 32% of the beads were detected as single beads (Supp. Fig. S1D). Therefore, anti-TfR and anti-MHCII were selected as control beads, and anti-BCR as activating beads, to assess the conjugate formation with mouse primary B cells. Cells and beads were incubated together for 15 minutes at 37 ºC and then analysed under the microscope. In addition, unbound cells were recovered and counted to determine the percentage of conjugates. Anti-BCR beads achieved the best conjugation efficiency such that around 70% of the cells attached to the beads (Fig. 1C), fitting well with the fraction of IgM^med/hi^ primary B cells (Mattila et al., 2013). Around 50% of the cells formed adhesions with the beads coated with anti-TfR and around 30% with anti-MHCII. We also tested beads coated with a combination of anti-TfR and anti-MHC-II but did not detect any improvement in conjugation. Therefore, from the tested control ligands, beads coated with anti-TfR antibodies performed the best and were selected as the negative control for further experiments. allowed us to proceed with these beads for subsequent protein isolation.

### B cell IS can be isolated using functionalized magnetic beads

Next, we validated the following steps of the pull-down Next, to verify our model system, we analysed the activation response of the primary B cells to the anti-BCR beads and the anti-TfR beads using microscopy and immunoblotting. Primary B cells were let to form conjugates with the coated beads in 1:1 ratio, fixed and stained with anti-phospho tyrosine (pY) antibodies and fluorescent phalloidin to assess the BCR signalling response and the IS formation, respectively. We clearly observed the two hallmarks of successful formation of the IS, the accumulation of F-actin and p Y signals at the site of the bead, in cells conjugated with anti-BCR beads, as measured by the intensity ratio (Fig. 2A). In contrast, the anti-TfR beads promoted no changes to the actin cytoskeleton nor to the pY signalling in the attached cells (Fig. 2A). Taken together, the anti-BCR coated beads successfully mimicked an APC and triggered IS formation, while the control anti-TfR beads induced cell:bead adhesion without BCR signalling or IS formation. This validation protocol: 1) chemical crosslinking of the proteins and 2) the sonication of the cells. These two parameters are of great importance, as crosslinking and sonication both need to be balanced in order to elute only the proteins in the cell adhesions without the bulk of the cells. Inefficient crosslinking or too strong sonication will result in great loss in recovered proteins, while on the other hand excessive crosslinking or insufficient sonication might lead to the pull-down of unspecific proteins and even nuclei.

**Figure 2.**
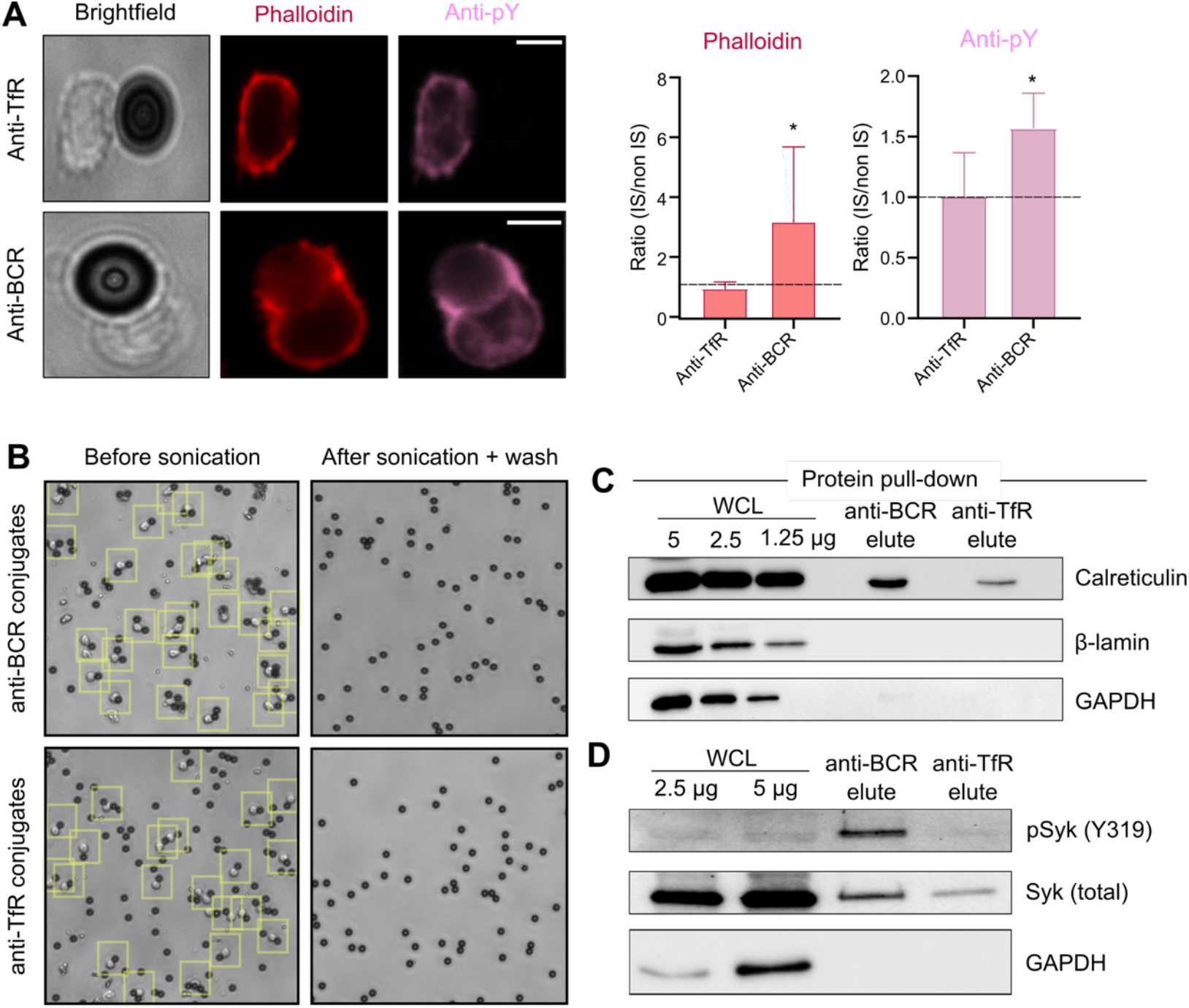
Validation of the specific formation and isolation of the IS. (A) Primary mouse B cells were let to form conjugates with the magnetic beads coated either with anti-BCR or anti-TfR antibodies, fixed, permeabilized, and stained for F-actin with fluorescent phalloidin and for tyrosine phosphorylation events using anti-p Y antibodies (left). The degree of BCR signalling was quantified by measuring the ratio of the mean signal intensity at the site of the bead as compared to the cell surface outside of the bead (on the right). Images were obtained with Spinning Disk confocal microscopy (*n* = 12). (B) Inverted brightfield microscopy was used to image cell-bead conjugates before and after sonication and washing of the cell debris. The results are representative of a set of 10 experiments. (C-D) 10 million cells and 10 million coated beads of each condition were processed with the IS isolation protocol and the bead bound IS proteins were eluted. (C) Specificity of the IS isolation was assessed by Western blotting with GAPDH (cytosolic protein), β-lamin (nucleus) and calreticulin (ER protein). WCL = whole cell lysate. (D) Successful extraction of the key BCR signaling kinase Syk in its phosphorylated form was evaluated by Western blotting, as compared to GAPDH as a control.

DTBP is a specific, cleavable and membrane-permeable crosslinker able to form stable amidine bonds with reactive amine groups (i.e., lysines and N-terminal amines). Different work groups have employed a range of DTBP concentrations (0.5 to 10 mM; (Byron et al., 2015; Jahan et al., 2018; Hammarén [MSc thesis], 2012), but the exact extent of DTBP activity in B cells is not described in the literature. Thus, we next evaluated the effect of DTBP in our system. To do so, we set up three small-scale samples for MS to test three different DTBP concentrations: 0.5 mM, 2.5 mM and 5 mM. The lowest concentration did not yield many proteins, suggesting that the crosslinking was not sufficient (data not shown). In this experiment, from samples with 2.5 mM DTBP, ≈500 total proteins were identified compared to 700 total proteins identified in pull-downs from 5.0 mM DTBP (data not shown). In order to be rather conservative in recovering the IS without excessive background from the non-specific parts of the cytosol, the subsequent experiments were performed with 2.5 mM DTBP. We also assessed the effect of 5-, 15- and 30-minutes incubation with DTBP on conjugate formation using anti-BCR coated beads, but found no difference between conditions (data not shown). Thus, the shortest time, 5 minutes, was selected.

Finally, we optimized the sonication to remove the cells from the beads leaving only the crosslinked proteins on the bead-adhesions for elution. Primary B cells isolated from mouse spleens were conjugated with anti-BCR beads (1:1) following the settings determined above and sonicated in multiple short rounds until essentially no cells were found attached to the beads (see Materials & methods for settings). After sonication, the samples were thoroughly washed to remove the debris and intact cells, yielding in very efficient removal of the cell bodies and leaving only the adhesion complexes on the beads (Fig. 2B). Next, we assessed the quality of the sonicated samples by immunoblotting. Proteins were eluted from the magnetic beads and blots were probed for calreticulin (ER marker), lamin (nucleus) and GAPDH (cytoplasm) to evaluate the presence of organelles not expected to pull down with the IS and, thus, potentially signs of unspecific background. Further suggesting high efficiency in sonication and subsequent washing steps to remove cell bodies, no lamin or GAPDH were found in the eluted samples (Fig. 2C). Interestingly, calreticulin was detected in the eluted samples suggesting trapping of some fragments of the ER in the adhesions, particularly in the IS (Fig. 2C). To evaluate the pull-down of BCR signalling proteins, we also assessed the presence of total and phosphorylated Syk, one of the first proximal tyrosine kinases phosphorylated upon antigen recognition, using Western blot. We observed a clear increase in the phospho-Syk to total Syk (pSyk/tSyk) ratio in those cells interacting with anti-BCR beads, compared to anti-TfR beads or unstimulated cells (Fig. 2D), further supporting the results shown in Figure 1A.

### Proteome of the immune synapse

With the above protocol set-up for isolation of the IS in primary mouse B cells, we proceeded to prepare samples for MS analysis. We conjugated primary B cells with anti-BCR and anti-TfR coated beads for 15 minutes, prior to proceeding to the IS extraction steps. Proteins halted to the beads were extensively washed to remove cell debris, contaminants and other non-crosslinked proteins. Then, they were lysed and trypsin digested for sequential mass spectrometry analysis employing an AP-MS/LC-MS approach. In total, we identified 2036 proteins (Fig. 3A, <1% false discovery rate [FDR]). After filtration of known contaminants and low confidence hits (characterized by having less than 1 unique peptide and a spectral count [MS count] lower than 5), we identified 1039 high confidence hits. A total of 661 proteins were exclusively found in anti-BCR samples, 365 proteins in both conditions, and only 13 proteins were exclusively found in the anti-TfR control samples. Among the proteins exclusively found in the control samples, we identified the TfR, confirming that we were able to specifically pull-down interacting proteins (Fig. 3A). Principal-component analysis (PCA) of all identified proteins was used to evaluate experimental fluctuation between biological replicates and conditions (Fig. 3B). The anti-TfR replicates were found very consistent while anti-BCR replicates showed more variance between each other. Overall, activated and non-activated conditions separated into two groups supporting their distinct nature (Fig. 3B-C).

**Figure 3.**
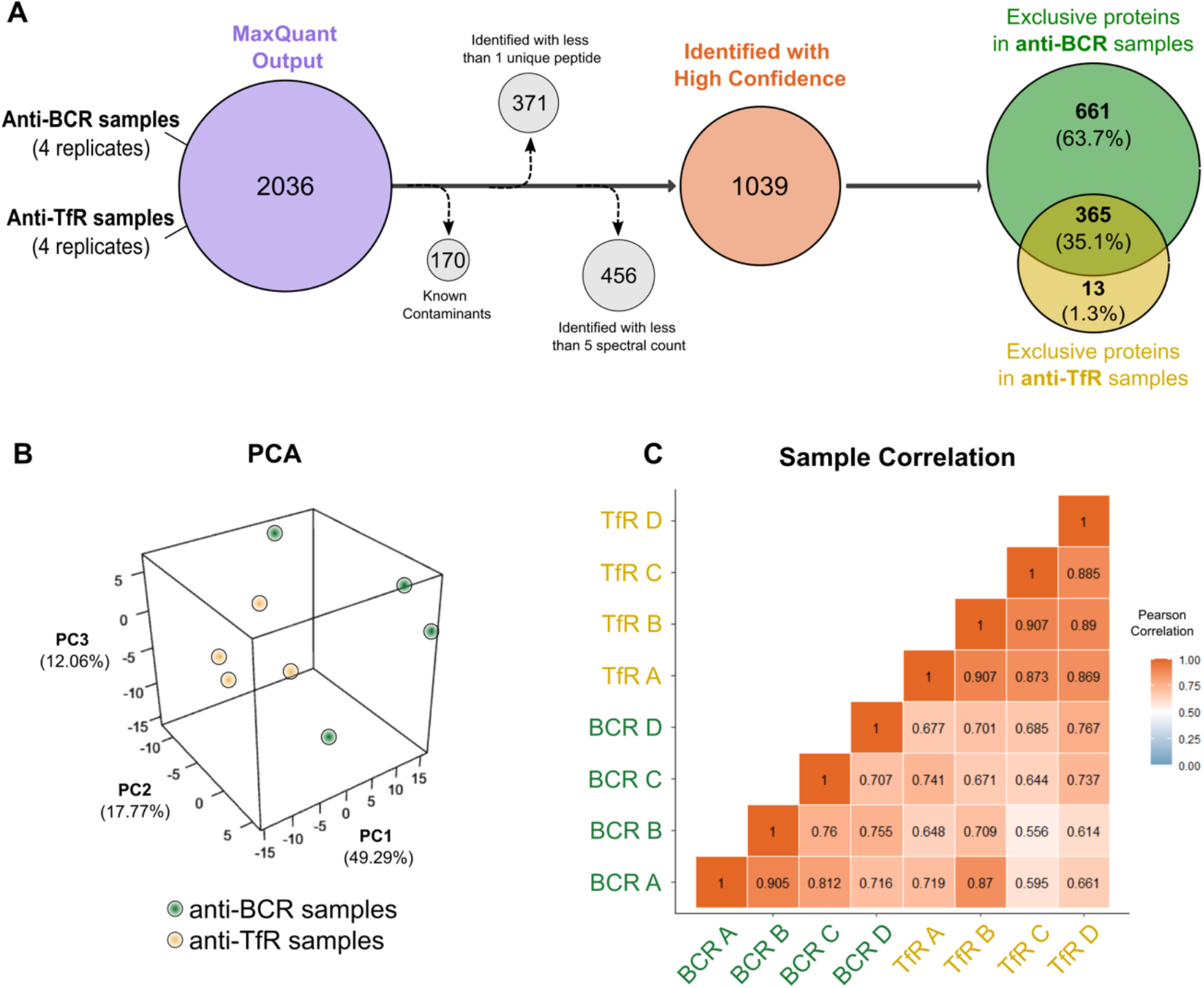
Mass spectrometry analysis of the isolated immune synapses. (A) Mouse primary B cells were let to form IS or control adhesions (anti-BCR and anti-TfR samples, respectively) for 15 prior to isolation. The extracted adhesion complexes were trypsin-digested and subjected for quantitative label-free MS analysis. 4 biological replicates were prepared. Total of 2036 proteins were identified by MaxQuant, and filtered as detailed in the Materials and Methods. The resulting high confidence proteins were further divided in three groups: exclusive present in either of the conditions or present in both conditions: 661 proteins were exclusively found in IS samples, 13 proteins in TfR samples, and 365 proteins in both conditions. (B) Principal component analysis of all the proteins identified with high confidence was done, after normalization using RLR normalization (Normalyz-erDE), in R environment. The three components accounting for 79% of the sample variability (PC1-3) are shown. (C) Pearson correlation was used to verify linearity and similarity relationship between biological replicates. TfR samples were highly correlated among replicates while IS samples featured more variability.

An important aspect to take into account in our data was that the anti-BCR (IS) samples typically featured higher overall protein content than the controls. The proteins bound on the beads were, in the end of the isolation protocol, eluted using Laemmli buffer with β-mercaptoethanol, allowing the extraction of the crosslinked IS proteins. The method of elution and the small yields of protein in total made the exact quantification of the samples highly challenging prior to the MS analyses. However, the protein amounts were recorded for each sample in the SDS-PAGE prior to trypsinization. We consistently saw 50% more extracted proteins in IS samples over the controls (Supp. Fig. S1E). This difference stems from the different conjugate formation rates between conditions and the lack of membrane extension over the beads in the control situation.

Next, we applied a differential enrichment (DE) analysis to the 365 proteins that were identified in high confidence in both sample types. DE analysis was performed in Perseus (Tyanova et al., 2016). The hierarchical clustering analysis highlighted various protein clusters with different enrichment profiles, with the largest ones showing clear enrichment in the IS over the control and contained various proteins previously linked to BCR activation such as Syk, Lyn, Vim, Ap2, Dock2, and Itsn2, among others (Fig. 4A). When we focused on the significantly enriched proteins, we also saw several proteins linked to the actin cytoskeleton such as Ezr and Dnm2 (Fig. 4B).

**Figure 4.**
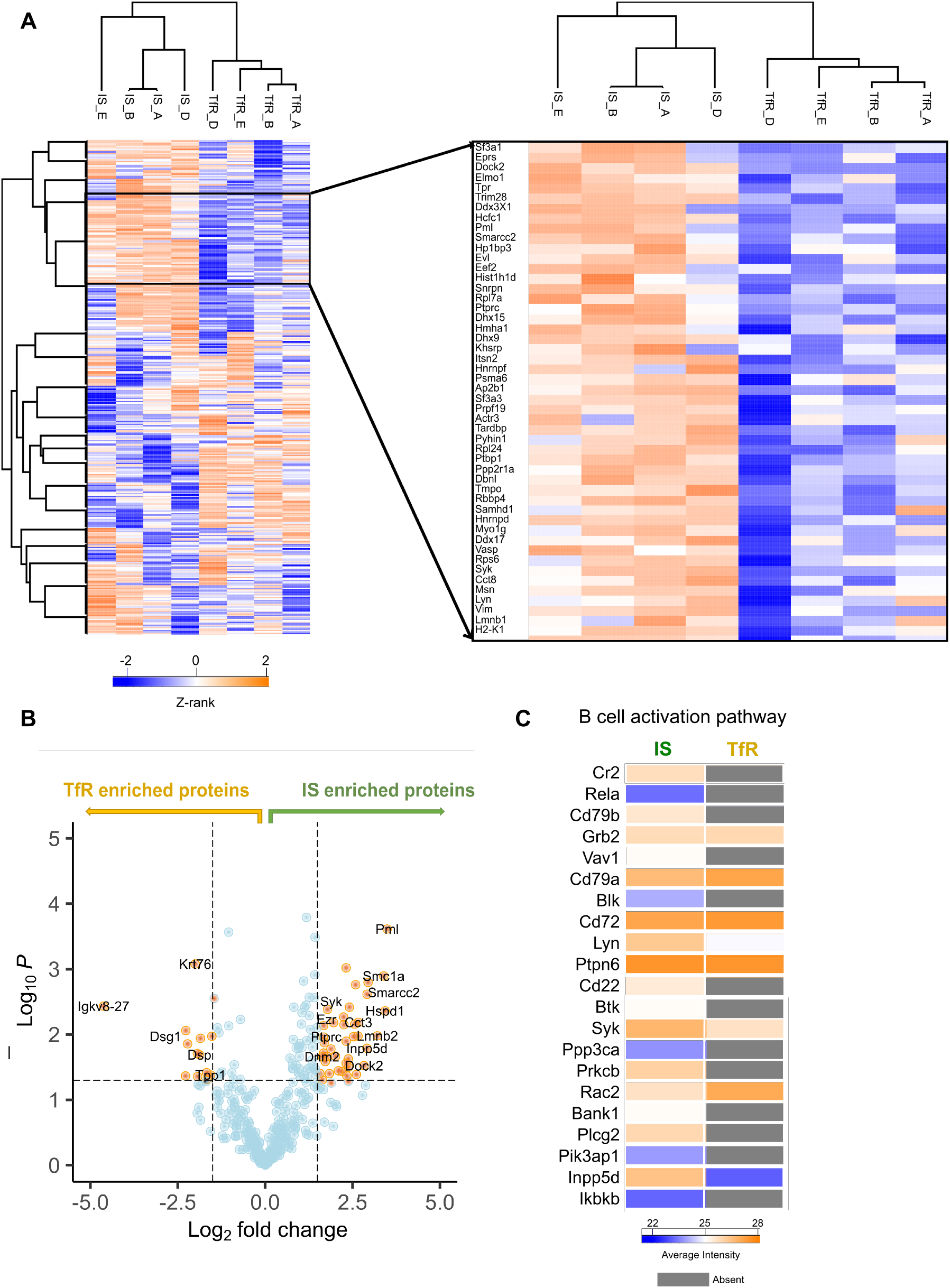
Enrichment analysis reveals protein accumulation to the IS. Clustering analysis of the 365 proteins identified in both isolated IS samples and the controls. (A) The heatmap showing hierarchical clustering was constructed by the z-rank score of all proteins shared between the conditions. Highly scored/enriched proteins have a z-rank score > 1 (orange), while modestly expressed have a z-rank score < −1 (blue). The cluster with the proteins most clearly enriched in the IS (zoom in; shown on the right side) contains various proteins previously described in B cell activation processes, such as Syk, Lyn, Dock2, Itsn2 and Psma6. (B) Differential enrichment analysis was carried out on the proteins identified in both conditions using Perseus. Differentially enriched proteins were identified using one-way ANOVA testing and p values were corrected for multiple testing using the method of Benjamini-Hochberg. The volcano plot depicts the Log_2_ fold changes of the protein intensity and the significance of the differential enrichment. Proteins found to be significantly differentially enriched (adjusted p-value ≤ 0.05 and a log_2_ fold change ≥ 1.5) in either sample are coloured in orange. (C) A heatmap of all the proteins identified in IS samples (exclusively or shared with the control) classified to the KEGG pathway “B cell activation”. Average protein intensities are plotted in each condition. Grey boxes indicate that protein is absent in that condition.

To further validate the specificity of our isolation method, we next addressed the behaviour of the known components of the BCR signalling cascade, namely proteins classified under the term B cell activation by KEGG pathway analysis, in the whole dataset (Fig. 4C). Majority of these proteins were exclusively identified in the IS samples, thus, they were absent form control adhesions. Similarly, most of these proteins were identified in relatively high intensities, and none of them in lower intensity in the IS as compared to the TfR adhesions (Fig. 4C). This data provided great confidence in our method to specifically isolate the IS proteins.

To obtain a general overview of the IS proteome we used the Over Representation Analysis (ORA) (Boyle et al., 2004) to determine which known biological functions or processes were over-represented among the proteins identified either exclusively or differentially enriched (log2 fold change ≥ 1.5 with an adjusted p-value ≤ 0.05) in the IS. This approach was applied using GO cellular components and KEGG pathways analysis (Fig. 5A and B). Interestingly, we found that the spliceosome and ribosomal complexes played a substantial role as more than 60 proteins hits belonged to these groups. The actual function of these proteins is not known in the IS, but the presence of these protein families has also been clearly detected APEX2based proximity proteomic screen (Awoniyi et al.) in B cells and in the proteomic study on T cell receptor (TCR) adhesions (De Wet et al., 2011). Expectedly, we also identified several modules involving the immune synapse, membrane structures, and actin protrusions linked to fundamental cell biological pathways such as regulation of the actin cytoskeleton and antigen uptake and processing (Fig. 5A and B). To illustrate selected molecular clusters in more detail, we generated interaction networks on proteins linked to BCR activation, actin cytoskeleton regulation, proteasome and endosomal traffic (Fig. 5C).

**Figure 5.**
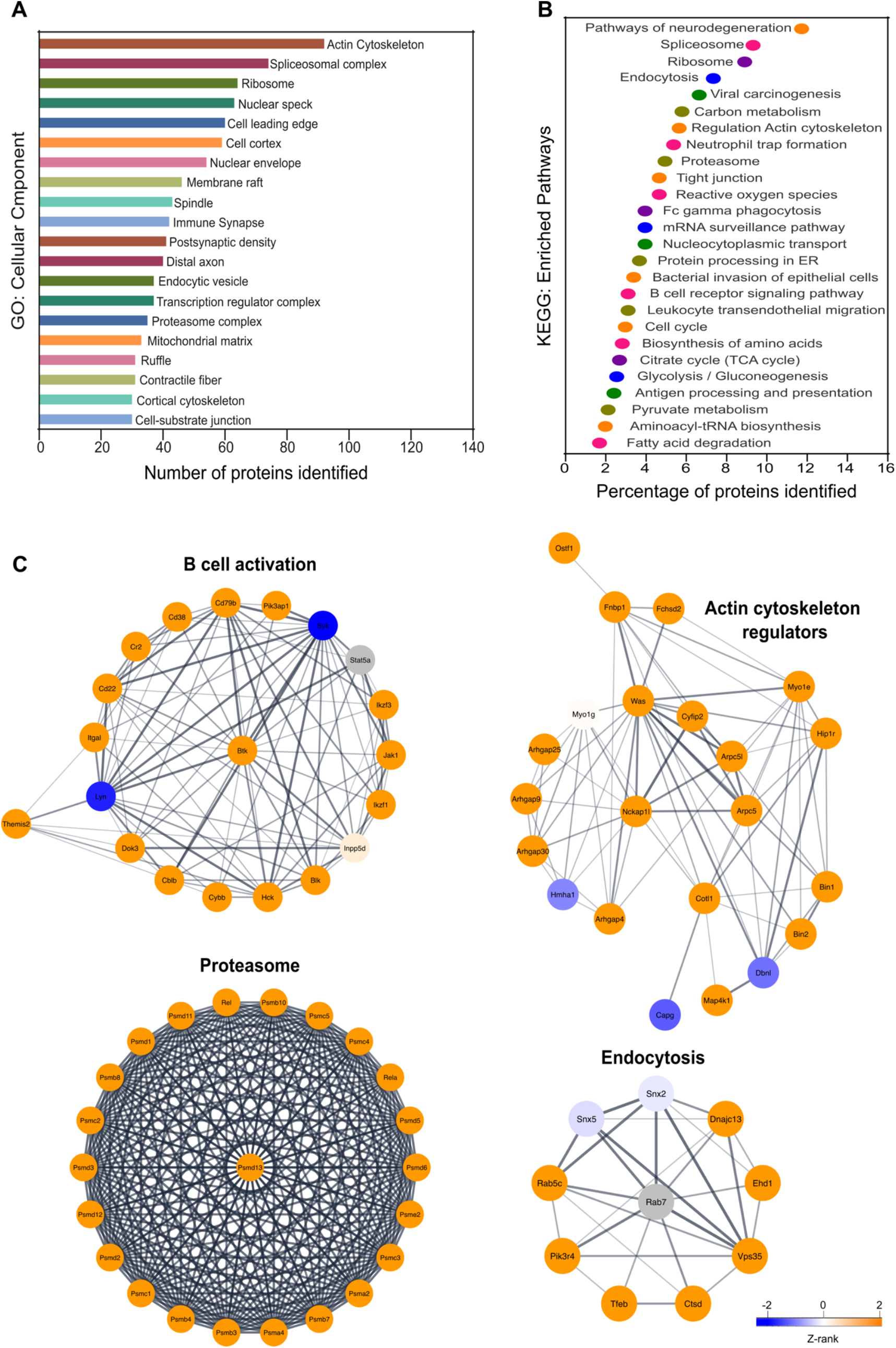
Proteins identified in the IS reveal both known and novel players participating in B cell activation. (A) An over-representation analysis based on Gene Ontology (GO) terms attributed to the cellular component of the 772 proteins identified in the isolated synapse (exclusive + shared with the control). GO terms that had ≥ 60% similarity were grouped and the one with lowest adjusted p-value is shown. (B) An over-representation analysis based on KEGG:pathways of the 772 proteins identified in the isolated synapses. Identified pathway terms that shared ≥ 60% similarity were grouped and the one with lowest adjusted p-value is shown. (C) High confidence proteins identified in IS samples were superimposed onto a composite protein-protein interaction (PPI) database of STRING. Full network was then filtered to provide edges with a high degree of confidence with a STRING score (>0.7). Filtered PPI network was then submitted to MCL clustering algorithm, an unsupervised cluster algorithm, with an inflation point of 2.5. Highest scored individual modules were annotated and further characterized in Cytoscape using three pathway databases. Nodes are coloured based on their z-rank score. Grey nodes are exclusively present proteins.

All together this data instils the confidence in our approach to detect the protein recruitment on the antigen-presenting substrates upon B cell activation.

## Discussion

The IS is a finetuned cell-cell interaction structure characteristic for lymphocytes and critical for their activation. Despite of the importance of this platform in antigen sensing, intracellular signalling, as well as endo- and exocytotic events, for instance, we have very little systems level information on the composition of the synapse. In this work, we have set-up a method for isolation the IS from mouse primary B cells for MS analysis. As a result, we report the proteomic content of the IS and compare it to cell adhesions formed by TfR to gain information on the specific molecules recruited by the BCR signalling. Our dataset reveals various known pathways as well as highlight potential new players for future studies.

The large-scale proteomic analyses have, in the recent years, unveiled various new features of cellular structures, facilitated by technical advancements both in the analysers as well as new molecular tools like spatial proteomics (Glass et al., 2020; Morgan & Tergaonkar, 2022). However, proteomic analyses on the IS has been lacking. Preservation of the membrane adhesions together with the linked cytosolic complexes challenges the method development. A bead-based system has been used for Jurkat T cells to capture of T cell receptor (TCR) membrane domains for proteomic analysis with anti-CD3-coated magnetic beads (De Wet et al., 2011; Harder & Kuhn, 2001) for the analysis. In this method, the separation of the TCR-linked adhesions, mimicking early stages of T cell IS, were separated from the rest of the cells with nitrogen cavitation. However, the efficiency of the isolation with this method was left unclear probably due to the potentially problematic background signal from other parts of the cells (Harder & Kuhn, 2001). In our IS isolation method, we took advantage of chemical crosslinking of the samples before removal of the cell bodies from the beads with sonication, similarly to the method developed to isolate integrin-based focal adhesions (Byron et al., 2015; Jones et al., 2015). Importantly, we were able to successfully apply our work-flow to isolate the IS from mouse primary B cells, which is a critical benefit over using cell lines. We detected only minimal residual GAPDH and lamin B, as markers for cytosolic content and nuclei (Fig. 2D). The sample replicates were well correlated with each other suggesting relatively low variability in the isolation procedures between the samples. The usage of DTBP cross-linker allowed retrieval of significantly higher amounts of proteins, suggesting to us that we were able to capture also the cytosolic protein networks in the IS. Notably, we also found that high concentrations of DTPB can induce selective protein phosphorylation, probably by stabilizing some kinase-based interactions or restricting phosphatase action. For this reason, great care would be required in applying a protein cross-linker to the samples that are destined for phospho-proteomic approaches.

To mimic antigen presenting surface, in our study, we utilized anti-BCR coated beads that have become a commonly utilized tool to study the formation of the IS (Yuseff et al., 2013). As a control, we chose anti-TfR antibodies to engage a well-studied cell surface receptor that has not been reported to induce any similar cellular changes than the engagement of the BCR, a feature that was also validated by us (Fig. 2A). From the different control molecules, we tested anti-TfR was among the best with its efficient coating-characteristics and efficiency in forming adhesions with B cells, although none of the control conditions tested yielded as high conjugation efficiency as surrogate antigen coated beads, again highlighting the prime efficiency of BCR engagement in B cell responses. Thus, in addition to the proteome data of the IS, we generated data on TfR adhesion proteome. TfR itself was among the 13 proteins only identified in control samples, providing trust in the quality of the control adhesions. For instance, enrichment of DSG1 and DSP in the TfR samples (Fig. 4B) propose for desmosome-like features of the TfR-mediated cell adhesion.

The proteomic data we obtained on the isolated IS unveiled a plethora of known and unknown players. The previously characterized BCR activation linked proteins were found largely exclusively in the IS samples or showed enrichment over the control samples, well validating the approach. The notable proteins networks found in the IS samples included, for instance, cytoskeletal regulators, vesicle transport mediators and proteasome complex, all with previously reported roles in B cell activation (De Wet et al., 2011; Ibañez-Vega et al., 2019; Kuokkanen et al., 2015; Tolar, 2017). Interestingly, we also found some large protein networks without previously characterized roles in the IS, like the spliceosome and ribosomal complexes. Together with the previous reports on the enrichment of these proteins at the sites of antigen receptor signalling (Awoniyi et al.; De Wet et al., 2011) these pathways emerge as attractive subjects for further studies.

## Acknowledgements

We thank Dr. Alexandre Carisey (St Jude’s Research Hospital, Memphis, USA) for his generous help with setting up the protocol. We also acknowledge Prof. Jordan Orange and Dr. Anna Meyer for their pioneering efforts in the immune synapse isolation in NK cells (Department of Pediatrics, Vagelos College of Physicians and Surgeons, Columbia University Irving Medical Center, New York, New York, USA). We thank Laura Grönfors for technical assistance. Microscopy and flow cytometry were performed at Turku Bioscience Cell Imaging and Cytometry (CIC), and proteomics at Turku Proteomics Facility, all supported by Turku Bioimaging and EuroBioimaging. The research infrastructures were provided by Biocenter Finland and InFLAMES.

## Author contributions

Conceptualisation: AFC, GJ, PKM. Formal Analysis: DMC, SHP. Funding Acquisition: PKM. Investigation: DMC, SHP. Methodology: DMC, SHP, LOA, AFC, GJ, PKM. Project Administration: SHP, PKM. Resources: PKM. Supervision: SHP, PKM. Visualisation: DMC, SHP. Writing - Original Draft Preparation: DMC, SHP, PKM. Writing - Review & Editing: DMC, SHP, LOA, AFC, GJ, PKM.

## Data availability

All data is available upon request.

## Materials and Methods

### Mice

Wild-type (WT) C57BL/6NCrl mice (Charles River Laboratories, Germany) were purchased from the University of Turku Central Animal Laboratory (UTU-CAL, Turku, Finland) and maintained under specific pathogen-free controlled conditions and in a temperature-controlled environment. Animals were housed at a maximum of 5 per cage in individually ventilated micro isolation cages and environmental enrichment was provided in the form of hiding igloos and nesting pads. Mice were fed regular chow ad libitum. Experiments were performed with mice between 8-12 weeks old. All animal experiments were approved by the Ethical Committee for Animal Experimentation in Finland and adhered to the rules and regulations of the Finnish Act on Animal Experimentation (62/2006; animal license numbers: 7574/04.10.07/2014, KEK/2014-1407-Mattila, 10727/2018).

### Mouse primary B cell isolation

A single-cell suspension of splenocytes was obtained by mechanical disaggregation of the spleen through a 70 μm cell strainer (#22363548, Thermo Fisher Scientific). Splenocytes were resuspended in B cell isolation buffer (Phosphate-buffered saline without Ca^+2^ and Mg^+2^ [PBS], 2% fetal calf serum [FCS], 1 mM EDTA) and isolated using a negative isolation kit (EasySep™ Mouse B Cell Isolation Kit, #19854, StemCell Technologies) according to the manufacturer’s instructions. Isolated B cells were let to recover in complete RPMI [cRPMI; RPMI 1640 with 2.05 mM L-glutamine supplemented with 10% FCS, 50 μM β-mercaptoethanol, 4 mM L-glutamine, 10 mM HEPES and 100 U/ml penicillin/streptomycin] in an incubator at +37 °C and 5% CO_2_ for at least 1 h before every experiment.

### Synapse Isolation

For bead preparation, 20 x 10^6^ Dynabeads™ M-450 Tosyl-activated (#14013, Thermo Fisher Scientific) were washed two times with Buffer 1 (0.1M sodium phosphate buffer, pH 7.4-8) and resuspended in Buffer 1 containing one of the following ligands: AffiniPure F(ab’)_2_ Fragment Goat Anti-Mouse IgM, μ chain specific (#115-006-020, Jackson ImmunoResearch) or rat anti-mouse CD71 (TfR) (#553264, BD Bioscience) (Cunha et al., *in press*). Beads were incubated in a shaker (1000 rpm) at RT for 90 min. After incubation, Buffer 1 with 1% BSA was added to the beads (final concentration of BSA 0.167%) and incubated for 16-24 hours at 37 ^º^C (1000 rpm). Beads were then washed three times with 20 mM HEPES and incubated for 2 hours at RT in 25 mM bis(sulfosuccinimidyl)suberate (BS3) (#21580, Thermo Fisher Scientific) diluted in 20 mM HEPES. After incubation, beads were washed with Buffer 3 (0.2 M Tris 0.1% BSA (w/v), pH 8.5) and incubated for 16-24 hours at RT (1000 rpm) to deactivate the remaining free tosyl groups on the beads. Beads were washed with Buffer 2 (PBS with 0.1% bovine serum albumin (BSA) and 2 mM EDTA, pH 7.4) and then stored in the same buffer for up to 1 month at 4^º^C. For cell - bead conjugation, cells were resuspended in Imaging Medium (RPMI 1640 Medium, no glutamine, no phenol red (#32404014, Thermo Fisher Scientific) supplemented with 1% FCS and 20 mM HEPES) and incubated in a 1:1 ratio with coated beads for 15 minutes at 37 ^º^C. After conjugation, Di-*tert*-butyl peroxide (DTBP) crosslinker (#20665, Thermo Fisher Scientific) was added for 5 minutes at 37 °C to a final concentration of 2.5 mM. Finally, DTBP was quenched by adding Tris-HCL 1M pH 8.5 to a final concentration of 150 mM for 5 minutes at RT. Quenched cell - bead conjugates were placed on ice and washed several times to remove unbound cells with cold Cytoskeletal Buffer ([CSK buffer]; 300 mM sucrose, 100 mM sodium chloride, 10 mM PIPES (pH 6.8), 3 mM magnesium chloride). Conjugates were resuspended in CSK buffer with 0.5% Triton X-100 and Pierce™ Protease Inhibitor Mini Tablets, EDTA-free (#A32955, Thermo Fisher Scientific) and sonicated in a BioRuptor sonicator (Bioruptor sonicator, Diagenode). Afterwards, conjugates were again washed 4-5 times with CSK buffer with 0.5% Triton X-100 to remove cell debris. Bead-bound proteins were then eluted in 2x Laemmli buffer (diluted from 4x Laemmli Sample Buffer, #161-0747, Bio-Rad) with betamercaptoethanol (#21985023, Thermo Fisher Scientific) by incubation at 70 ^º^C for 30 min followed by a second incubation at 95 ^º^C for 10 min. Beads were discarded, and eluted proteins were used immediately or stored at - 20≡C for up to 1 week.

### Flow Cytometry

To verify the antibody coating, beads coated with anti-IgM or anti-CD71 were stained with fluorescently labelled secondary antibodies (donkey anti-goat IgG Alexa Fluor 488 (#A11055, Thermo Fisher Scientific) or donkey anti-rat IgG Alexa Fluor 488 (712-545-153, Jackson ImmunoResearch), respectively). For detection of cell:bead conjugates, conjugates were fixed in 4% PFA, washed, and stained with Purified Rat Anti-Mouse CD16/CD32 (Mouse BD Fc Block™) (#553142, BD Biosciences), Donkey anti-Goat IgG Alexa Fluor 488 (#A11055, Thermo Fisher Scientific) and DAPI (#D1306, Thermo Fisher Scientific) for 30 minutes. Data were acquired using a BD LSR Fortessa analyser equipped with four lasers (405, 488, 561 and 640 nm) and analysed using FlowJo v10 (Tree Star).

### Brightfield and fluorescence microscopy

To evaluate the formation of conjugates and the effect of the sonication, samples were diluted 1:20 in PBS and analysed using an inverted microscope EVOS fl, with an illumination system based on LED cubes (Thermo Fischer). The images were captured using a 40x AMG PlanFluor objective and 4 different filter sets: DAPI, GFP, RFP, and transmitted light.

### Spinning disk confocal microscopy

Conjugates were developed from a 1:1 cell - bead ratio. Unbound cells were removed and conjugates were fixed with PFA 4% at RT for 20 minutes. Conjugates were further stained with fluorescent phalloidin (Alexa Fluor 555) and anti-pY antibody to assess the degree of BCR activation with each set of coated beads. Images were acquired using a 3i CSU-W1 spinning disk equipped with 405, 488, 561 and 640 nm laser lines and 510-540, 580-654 and 672-712 nm filters and 63× Zeiss Plan-Apochromat objective. SlideBook6 (Intelligent Imaging Innovations Incorporation) software was used for image acquisition.

### Immunoblotting

For the analysis of BCR signalling and protein recruitment (lamin, GAPDH and calreticulin), eluted proteins were run on 10% polyacrylamide gels and transferred to polyvinylidene fluoride (PVDF) membranes (Trans-Blot Turbo Transfer System, BioRad). Membranes were blocked with 5% milk in Tris-buffered saline (TBS, pH ~7.4) for 1 h and incubated with primary antibodies in 5% BSA in TBST (TBS, 0.05% Tween-20) O/N at 4°C. Secondary antibody incubations were done for 1h at RT in 5% milk in TBST for horseradish peroxidase (HRP)-conjugated antibodies. Washing steps were done in 10 ml of TBST for 5 × 5 min. Membranes were scanned with ChemiDoc MP Imaging System (Bio-Rad) after the addition of Immobilon Western Chemiluminescent HRP Substrate (WBKLS0500, Millipore) and incubation of 5 minutes. Images were background subtracted, and the raw integrated densities for each band were measured in ImageJ.

### In-gel digestion

Eluted samples were run for about 1 cm on Any kD™ Mini-PROTEAN^®^ TGX™ Precast Protein Gels (#4569034, Bio-Rad), and the gel was stained with Pierce™ Zinc Reversible Stain Kit (#24582, Thermo Fisher Scientific) for protein quantification. Gel pieces were cut from the gel (3 pieces per lane) and distained in Tris-glycine pH 8.0 (25 mM Tris, 192 mM glycine), followed by three washes with MQ water for 10 minutes. Gel pieces were then further cut into four pieces and processed as previously described with the following modifications (Shevchenko et al., 2007). Gel pieces were shrunk by covering the gel with 100% Pierce™ Acetonitrile (ACN), LC-MS Grade (#51101, Thermo Fisher Scientific) for 5-10 minutes. Then, gel pieces were rehydrated for 30 minutes at 56 ^º^C with 20 mM Dithiothreitol (DTT) (#20291, Thermo Fisher Scientific) and dehydrated again with 100% ACN. Then, gel pieces were rehydrated in 55 mM iodoacetamide (IAA) (CAS:144-48-9, Sigma-Aldrich) for 20 min in the dark RT and washed with 100 mM ammonium bicarbonate (NH_4_HCO_3_) (09830-500G, Sigma-Aldrich) and dehydrated as above. Afterwards, 0.02 μg/μl Sequencing Grade Modified Trypsin (v511A, Promega) was added to the gel pieces and let to absorb for 20 minutes on ice. When all trypsin solution was absorbed, 40 mM NH_4_HCO_3_ / 10% ACN was added to cover the gel pieces and incubated for 18 hours at 37 ^º^C. After incubation, 100% ACN was added and incubated for 15 min at 37 ^º^C. Finally, the supernatant was collected into a clean Eppendorf tube, and the extraction was repeated with 50 % ACN / 5 % Formic Acid (HCOOH) (LC-MS Grade, #85178, Thermo Fisher Scientific). The solution was dried in vacuum centrifugation for 2 hours 30 minutes (maximum vacuum pressure up to 5.1; heating time of 20 minutes at 45 ^º^C

### Mass spectrometry analysis

Data was collected by LC - electrospray ionisation (ESI) - MS/MS using a nanoflow HPLC system (Easy-nLC1200, Thermo Fisher Scientific) coupled to the Orbitrap Fusion Lumos mass spectrometer (Thermo Fisher Scientific) equipped with a nano-electrospray ionisation source. Peptides were first loaded on a trapping column and subsequently separated inline on a 15 cm C18 column (75 μm x 15 cm, ReproSil-Pur 3 μm 120 Å C18-AQ, Dr. Maisch HPLC GmbH, Ammerbuch-Entringen, Germany). The mobile phases consisted of water with 0.1% HCOOH (solvent A) and ACN/water (80:20 (v/v)) with 0.1% HCOOH (solvent B). A 60 min chromatography method was used: from 5% to 21% of solvent B in 28 min, to 36% of solvent B in 22 min, from 36% to 100% of solvent B in 5 min, followed by a wash at 100% of solvent B for 5 min. MS data were acquired automatically using Thermo Xcalibur 4.1 software (Thermo Fisher Scientific). A data-dependent acquisition method (DDA) consisting of an Orbitrap MS survey scan of mass range 350-1750 m/z followed by HCD fragmentation was used.

### Protein identification and differential enrichment analysis

Raw files were processed with MaxQuant software version 1.6.17.0 (Cox et al., 2008) with the first search peptide tolerance set to 20ppm. Trypsin was set as the digestion enzyme, carbamidomethyl specified as fixed modification, and oxidation, acetylation (Protein N-term), DTBP + alkylation (K) and DTBP + alkylation (Protein N-term) were specified as variable modifications. MS/MS spectra were searched against the mouse Swiss-Prot database (17, 132 protein entries; downloaded in01/2021). Peptide and protein false discovery rate were set to 0.01. MaxLFQ and match between runs were enabled.

Proteins identified through MaxQuant were subjected to several filtering steps: potential contaminants, proteins only identified by site, reverse hits, proteins with less than 1 unique peptide and proteins with a spectral count lower than 5. Additionally, proteins with missing values in 3-4 conditions in each conditions were filtered out. These high confidence proteins were further divided in two groups: exclusive present in either condition or differently enriched in either condition.

High confidence proteins were then evaluated in a quantitative and qualitative manner using several normalization methods through NormalyzerDE (Willforss et al., 2019). RLR normalization was deemed the best normalization method (data not shown). For IS enrichment analysis, one-way ANOVA was used to identify DE proteins, and p values were corrected for multiple testing using the method of Benjamini-Hochberg. Several missing value imputation methods were performed and evaluated and ultimately, deciding on using k-Nearest Neighbor (kNN) Imputation, using the 15 closest neighbours as adjustment. Differential expression analysis was done using Perseus (Tyanova et al., 2016). Principal-component analysis and hierarchical clustering were performed using R (v.4.2.2).

### Protein Network Analysis

High confidence proteins exclusively present or enriched in IS samples were superimposed onto a composite protein-protein interaction (PPI) database of STRING (http://string-db.org/) (Szklarczyk et al., 2019). String database provides an enormous number of PPIs since it collects information from all sources, edges considered were edges with a high degree of confidence, as determined by the STRING score (>0.7). Filtered PPI network was then submitted to MCL clustering algorithm, an unsupervised cluster algorithm, with an inflation point of 2.5. Obtained individual modules were annotated and further characterized in Cytoscape (Doncheva et al., 2019) using three pathway databases (Gene Ontology, KEGG and Reactome).

### Illustrations and statistics

Graphs were created in GraphPad Prism 8, and illustrations were created with BioRender. Figure formatting was done on Inkscape 1.0. The Word template for bioRxiv submission was downloaded from chrelli.github.io.

---

**Figure S1:**
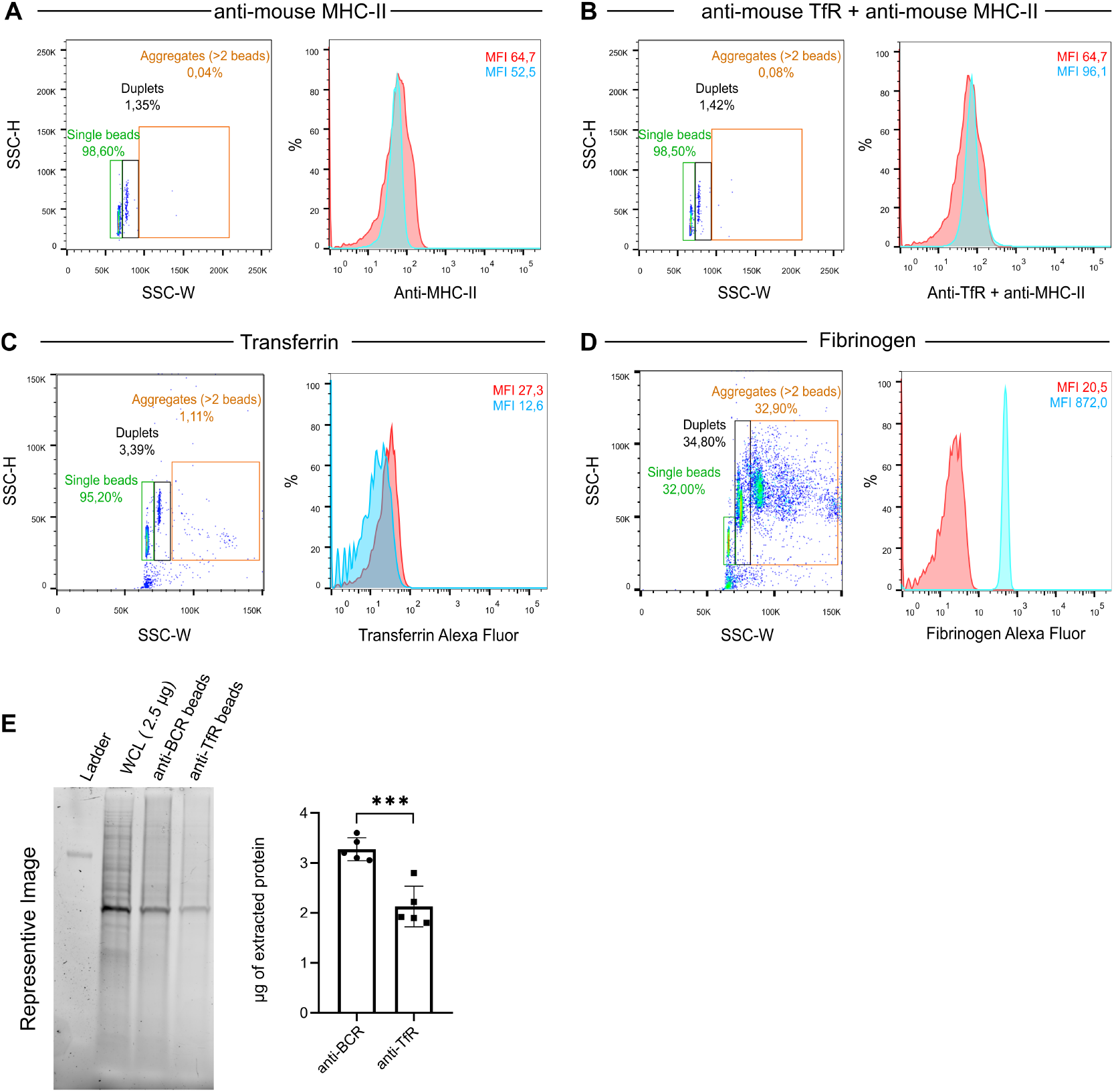
Screening of control ligands for non-activating cell adhesions. (A-D) Coating efficiency of the beads with the control ligands [(A) anti-MHC-II, (B) combination of anti-TfR and anti-MHC-II, (C) transferrin, and (D) fibrinogen] was determined using flow cytometry. Single beads (green), duplets (black) and aggregates (orange) were gated separately (on the left) and the fluorescence intensity of the single beads was analysed (on the right). Red histogram: uncoated beads. Blue histogram: coated beads. (E) 10 million cells and 10 million coated beads of each condition were let to form conjugates for 15 minutes, processed with the IS isolation protocol and the bead-bound IS proteins were eluted. Samples were loaded on an SDS-PAGE gel and the total amount of protein was quantified using Zinc staining. A known amount of whole cell lysate (WCL) was used as a control. A representative blot is shown on the left, and quantification from n = 5 experiments is shown on the right. *** P < 0.0001 (unpaired Student’s t-test).

